# Comparative analysis of total alpha-synuclein (αSYN) immunoassays reveals that they do not capture the diversity of modified αSYN proteoforms

**DOI:** 10.1101/2022.03.11.483980

**Authors:** Lara Petricca, Nour Chiki, Layane Hanna-El-Daher, Lorène Aeschbach, Ritwik Burai, Erik Stoops, Mohamed-Bilal Fares, Hilal A. Lashuel

**Author notes:** Corresponding Author, Laboratory of Molecular and Chemical Biology of Neurodegeneration, EPFL SV BMI LMNN, AI 2151 (Bâtiment AI), Station 19, CH-1015 Lausanne.

## Abstract

**Background:** The development of therapeutics for Parkinson’s disease (PD) requires the establishment of biomarker assays to enable stratifying patients, monitoring disease progression and assessing target engagement. Attempts to develop diagnostic assays based on detecting levels of the α-synuclein (αSYN) protein, a central player in the pathogenesis of PD, have yielded inconsistent results.

**Objective:** To determine whether the three commercial kits that have been extensively used for total αSYN quantification in human biological fluids (from Euroimmun, MSD, and Biolegend) are capable of capturing the diversity and complexity of relevant αSYN proteoforms.

**Methods:** We investigated and compared the ability of the different assays to detect the diversity of αSYN proteoform using a library of αSYN proteins that compromise the majority of disease-relevant αSYN variants and post-translational modification.

**Results:** Our findings showed that none of the three tested immunoassays accurately capture the totality of relevant αSYN species and are unable to recognize most disease-associated C-terminally truncated variants of αSYN. Moreover, several N-terminal truncations and phosphorylation/nitration differentially modify the level of αSYN detection and recovery by different immunoassays, and a CSF matrix effect was observed for most of the αSYN proteoforms analyzed by the three immunoassays.

**Conclusions:** Our results showed that these immunoassays do not capture the totality of the relevant αSYN species and therefore may not be appropriate tools to provide an accurate measure of total αSYN levels in samples containing modified forms of the protein. This highlights the need for next-generation αSYN immunoassays that capture the diversity of αSYN proteoforms.

## Introduction

Parkinson’s disease (PD) is a neurodegenerative disorder whose exact cause is still unknown and for which no therapy is available today. The number of patients suffering from PD is projected to increase from 6.1 million (reported in 2016 [1]) to 9.3 million by 2030 [2], rendering PD one of the fastest growing neurological disorders worldwide. Dopaminergic therapies represent the gold standard for symptom management of PD, but intervention to modify and slow down the clinical course of the disease remains a major unmet need, as many disease-modifying therapies targeting different pathogenic pathways in PD have recently failed to meet their primary endpoints in clinical trials [3–7]. Among the different reasons underlying these recent failures is the clinical heterogeneity of PD and the lack of sensitive biomarkers and assays that enable early diagnosis of PD, patient stratification, assessment of target engagement, and monitoring of disease progression.

Currently, the diagnosis of PD is mainly based on the medical history of the patients, a review of clinical symptoms, and neurological and physical examination following the International Parkinson and Movement Disorder criteria [8]. Given that the first symptoms of PD are also common to other neurodegenerative movement disorders or medical conditions, PD is often misdiagnosed. In fact, accurate clinical diagnosis is often made only when ≈ 70% of the dopaminergic neurons have already degenerated, a very late stage of disease progression [9]. This, combined with the fact that the disease starts 10-15 years before the manifestation of the characteristic clinical symptoms of the disease, underscores the critical importance of developing biomarker assays not only to support accurate and early diagnosis but also to monitor disease progression, enable patient stratification and interrogate the efficacy of therapeutics in the clinic.

Over the past 20 years, the protein α-synuclein (αSYN) has attracted increasing interest as a central player in the pathogenesis of PD and as one of the primary therapeutic targets for the treatment of PD. This was mainly following the seminal discovery that mutations in the SNCA gene encoding αSYN cause familial PD [10,11] and that aggregated and posttranslationally modified (PTM) forms of αSYN, including phosphorylated, nitrated, ubiquitinated, and truncated species, are key components of Lewy bodies (LB) and Lewy neurites, the main neuropathological hallmark of PD and other synucleinopathies [12–14], [15]. Furthermore, the finding that different αSYN proteoforms could be detected in biological fluids (cerebrospinal fluid (CSF), plasma and red blood cells (RBC), and saliva) [16][17] [18–29] as well as in peripheral tissues (skin, intestinal mucosa, submandibular gland) [30–33] stimulated interest in αSYN as a potential disease biomarker for the early diagnosis of PD and for differentiation between PD and other synucleinopathies [25,34–36].

Several αSYN-targeting therapeutic approaches are being pursued in academia and industry, underscoring the need for sensitive αSYN assays to assess target engagement and therapeutic efficacy. Nevertheless, accurate detection and quantification of total αSYN and pathologically relevant variants continues to represent a major challenge, as demonstrated by the marked discrepancies in αSYN levels reported by different studies and laboratories. Whereas several independent groups have reported lower total αSYN levels in the CSF of PD patients compared to control subjects [20,22–26,28,29], other groups have found no significant differences [27,32]. Similarly, attempts to use total CSF αSYN levels to distinguish among different types of synucleinopathies, including PD, MSA, DLB, and PSP, have yielded inconsistent results [25,34–36]. Furthermore, large variations and discrepancies have been reported in the quantified levels of total αSYN using different assays. For instance, in a recent comparative study using the same human CSF samples [37], although four of the most commonly used commercial immunoassays for total αSYN detection revealed similar overall correlations across cohorts of patients and controls, large variations in the absolute concentrations of total αSYN were quantified. Indeed, the Euroimmun (Euroimmun AG, Lübeck, Germany, developed by ADx Neurosciences, Gent, Belgium) and BioLegend Legendmax (San Diego, California, USA) immunoassays gave much higher concentrations than the MSD U-PLEX^®^ (Meso Scale Discovery, Rockville, Maryland) and Roche Elecsys^®^ (Roche Diagnostics, Penzberg, Germany) assays (334–3547 pg/mL, compared to 35.1–607 pg/mL).

Several factors have been proposed to explain such variations in total αSYN detection, including 1) the clinical heterogeneity of patients; 2) differences in sample-handling procedures; 3) differences in the sensitivity of the assay platforms; and 4) the use of different protein standards in absence of a certified reference material. In addition, we hypothesized that the antibodies deployed in most of these assays might not capture the full diversity of relevant αSYN proteoforms in biological samples. Indeed, increasing evidence shows that αSYN in brain LBs, peripheral tissues, and biological fluids exists as a mixture of different PTM forms and conformations, including different phosphorylated, truncated, nitrated, and ubiquitinated forms of αSYN [38,39][40] [41]. Moreover, the levels of several modified αSYN proteoforms have been reported to be higher in the biological fluids and tissues of PD patients than in those of healthy controls. Furthermore, the distribution of αSYN proteoforms was recently shown to differ between the soluble and insoluble states of the protein and between brain tissues and peripheral tissues (e.g., appendix [33]). Altogether, these observations suggest that assays that do not account for this diversity may not accurately report on the total levels of αSYN.

Notably, most of the antibodies deployed in commercial immunoassays for total αSYN target the C-terminal domain of αSYN (Figure 1), which is known to harbor the majority of αSYN PTMs, including phosphorylation nitration and C-terminal truncations [38,39]. It is possible that these PTMs could either eliminate or mask the epitope of these antibodies or interfere with antibody binding to αSYN resulting in a biased analyte concentration. The C-terminus of the αSYN protein is the most immunogenic region and results in high affinity antibodies and thus better assay performance whereas αSYN N-terminal antibodies have in general a lower affinity resulting in lower sensitivity assays. That is most likely the reason for selection of the antibody pairs in the commercial assays. In addition, the great majority of αSYN immunoassays were developed based on the use of a single protein standard (full-length αSYN), which may have led to the downstream development of assays that do not necessarily capture the diversity of αSYN proteoforms.

**Figure 1.**
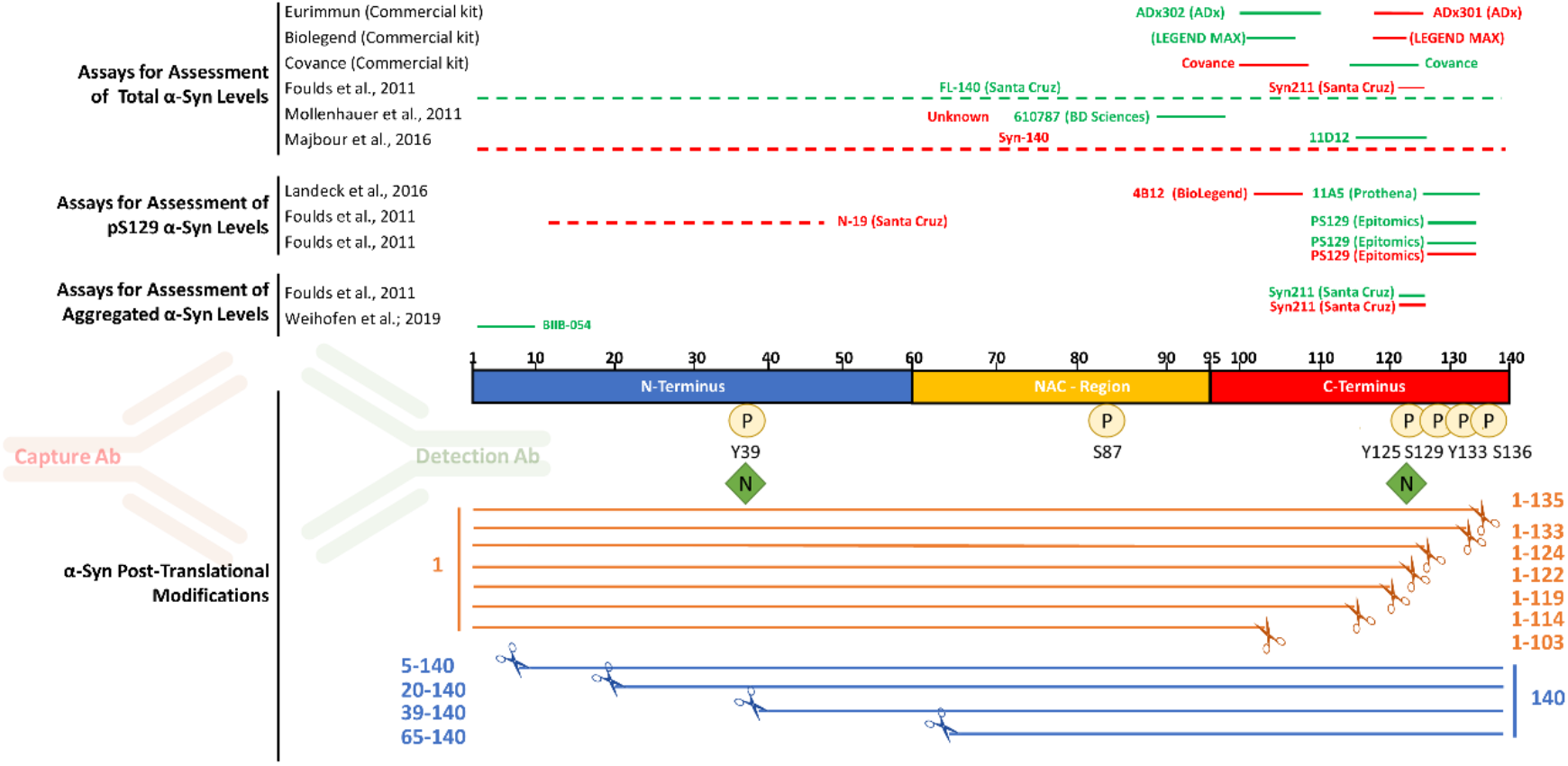
Most antibodies used in immunoassays target the C-terminal posttranslational modified region of αSYN. Schematic depiction of the main αSYN PTMs (phosphorylation, nitration, and truncation). Red and green bars above the sequence of αSYN show the epitopes of capture and detection antibodies, respectively, used in immunoassays to measure total αSYN, pS129 αSYN, and aggregated αSYN levels in human biological specimens.

In this study, we sought to test this hypothesis by systematically investigating whether the three commercial kits that have been extensively used for total αSYN quantification in human biological fluids (from Euroimmun, MSD, and Biolegend) are capable of detecting the diversity and complexity of αSYN proteoforms. For this purpose, we capitalized on our ability to reconstitute the diversity of αSYN PTM species under controlled conditions [42–45],[46],[47]. Specifically, we assessed the ability of the different assays to detect different αSYN proteoforms harboring N- and C-terminal truncations and other site-specific PTMs, such as phosphorylation and nitration at different residues. Altogether, our findings show that the three analyzed immunoassays do not capture the totality of the relevant αSYN species. More specifically, the three assays fail to detect the most common disease- and pathology-associated truncated αSYN species. Furthermore, N-terminal truncations and PTMs (phosphorylation/nitration) differentially modify the level of αSYN detection and recovery by different immunoassays. The implications for developing assays that allow the accurate quantification of total αSYN levels and the assessment of αSYN as a diagnostic biomarker for PD and other synucleinopathies are discussed. These findings suggest that these assays are not appropriate tools to provide an accurate estimate of total αSYN levels in biological samples that are likely to contain modified forms of the protein.

## Materials and methods

### Preparation and characterization of the αSYN protein library

All proteins except pS129 αSYN were generated at the Lashuel lab using recombinant protein expression, protein semisynthesis, or previously described total chemical synthesis strategies [42–45], [48], [46],[47]. The purity of all proteins was independently validated by the Lashuel lab and ND Biosciences by multiple techniques, including Coomassie staining, mass spectrometry and UPLC, which together established the purity and integrity of the generated proteins. Lyophilized samples were resuspended in 1x Tris-buffered saline and passed through a 100 kDa filter. Human alpha-synuclein phospho-Ser129 protein (Cat. No. RP-004) was purchased from Proteos (Proteos, Inc., USA) and resuspended according to the manufacturer’s instructions. All proteins were quantified by amino acid analysis to establish accurate protein concentrations (Functional Genomics Center Zurich, ETH Zürich).

### Human CSF samples

Commercial pooled human CSF was acquired from Innovative Research Inc. (Cat. No. IRHUCSF5ML) and stored in aliquots frozen at −20 °C until use.

### Assessment of the in-house αSYN protein library using total αSYN commercial immunoassay kits

The <u>LEGEND MAXtrademark Human *α*-Synuclein ELISA Kit (BioLegend; Cat. No. 844101)</u> was kindly supplied by Biolegend. The assay was performed according to the manufacturer’s instructions. In brief, full dilution curves of the 10 proteins with the same dynamic range as the standard curve of the kit-standard (6.1–1500 pg/ml) were obtained by serial dilutions in assay buffer. Calibration curve points and protein samples were loaded in duplicate. Luminescence was measured using a FLUOstar plate reader (BMG Labtech). The results were fitted to a 4-parameter sigmoid curve and plotted against the kit standard curve. Further details can be found in Figure S8.

Spike recovery experiments were performed by spiking high (900 pg/ml), medium (300 pg/ml) and low (50 pg/ml) amounts of each protein (according to the dynamic range of the kit’s standard curve) into assay buffer and into commercial human CSF (hCSF). hCSF samples were diluted 1:20 according to the manufacturer’s instructions. Recovery was assessed as the percentage of the back-calculated values of the spiked samples (subtracted of the unspiked sample) with respect to the nominal spiked protein amount. The high, medium and low recovery percentage were averaged.

<u>U-PLEX Plus Human Alpha-Synuclein Kit (Mesoscale Discovery; Cat. No. K151 WKP-2)</u> was acquired from Meso Scale Diagnostics. The assay was performed according to the manufacturer’s instructions. In brief, full dilution curves of the 10 proteins with the same dynamic range as the standard curve of the kit-standard (2.44–10000 pg/ml) were obtained by serial dilutions in assay buffer. Calibration curve points and protein samples were loaded in duplicate. Plates were read using a MESO QuickPlex SQ 120 (Mesoscale Discovery). The results were fitted to a 4-parameter sigmoid curve and plotted against the kit standard curve. Further details can be found in Figure S8.

Spike recovery experiments were performed by spiking high (7000 pg/ml), medium (1000 pg/ml) and low (100 pg/ml) amounts of each protein (according to the dynamic range of the kit’s standard curve) into assay buffer and into commercial human CSF (hCSF). hCSF samples were diluted 1:20 according to the manufacturer’s instructions. Recovery was assessed as the percetage of the back-calculated values of the spiked samples (subtracted of the unspiked sample) with respect to the nominal spiked protein amount. The high, medium and low recovery percentage were averaged.

<u>Euroimmun Alpha-Synuclein ELISA (Euroimmun; Cat. No. EQ 6545-9601-L)</u> was kindly supplied by ADx (ADx NeuroSciences NV, Gent, Belgium) and detects the C-terminus of α-synuclein [49] [50]. The assay was performed according to the manufacturer’s instructions. In brief, full dilution curves of the 10 proteins with the same dynamic range as the standard curve of the kit standard (150–5988 pg/ml) were obtained by serial dilutions in assay buffer. Calibration curve points and protein samples were loaded in duplicate. Absorbance was measured using a VarioSkan Lux plate reader (Thermo Scientific). The results were fitted to a 4-parameter sigmoid curve and plotted against the kit standard curve. Further details can be found in Figure S8.

Spike recovery experiments were performed by spiking high (4000 pg/ml), medium (1535 pg/ml) and low (348 pg/ml) amounts of each protein (according to the dynamic range of the kit’s standard curve) into assay buffer and into commercial human CSF (hCSF). hCSF samples were diluted 1:10. Recovery was assessed as the percentage of the back-calculated values of the spiked samples (with the unspiked sample subtracted) with respect to the nominal spiked protein amount. High, medium and low recovery % were averaged.

### Data analysis

Data analysis and creation of the corresponding graphs were performed using GraphPad Prism 9 (GraphPad Software, Inc., San Diego, CA).

## Results

### Generation and characterization of protein standards

To determine the capability of the three commercial immunoassays (from Euroimmun, BioLegend and MSD) to capture the diversity of αSYN species, we first assembled a library of 15 highly pure αSYN proteins bearing the most commonly occurring PTMs as previously described [42–45] (Figure 2). The purity and integrity of all the proteins were validated by multiple orthogonal techniques, including Coomassie staining, mass spectrometry and UPLC (Supplemental Figure 1). Furthermore, to ensure that any slight differences in assay detection were attributable to different affinities of deployed antibodies toward different proteins and not to differences in protein loading onto assay plates, the concentration of all proteins was determined by amino acid analysis (AAA), which is the gold standard allowing the most precise determination of protein concentrations [51] [52].

**Figure 2.**
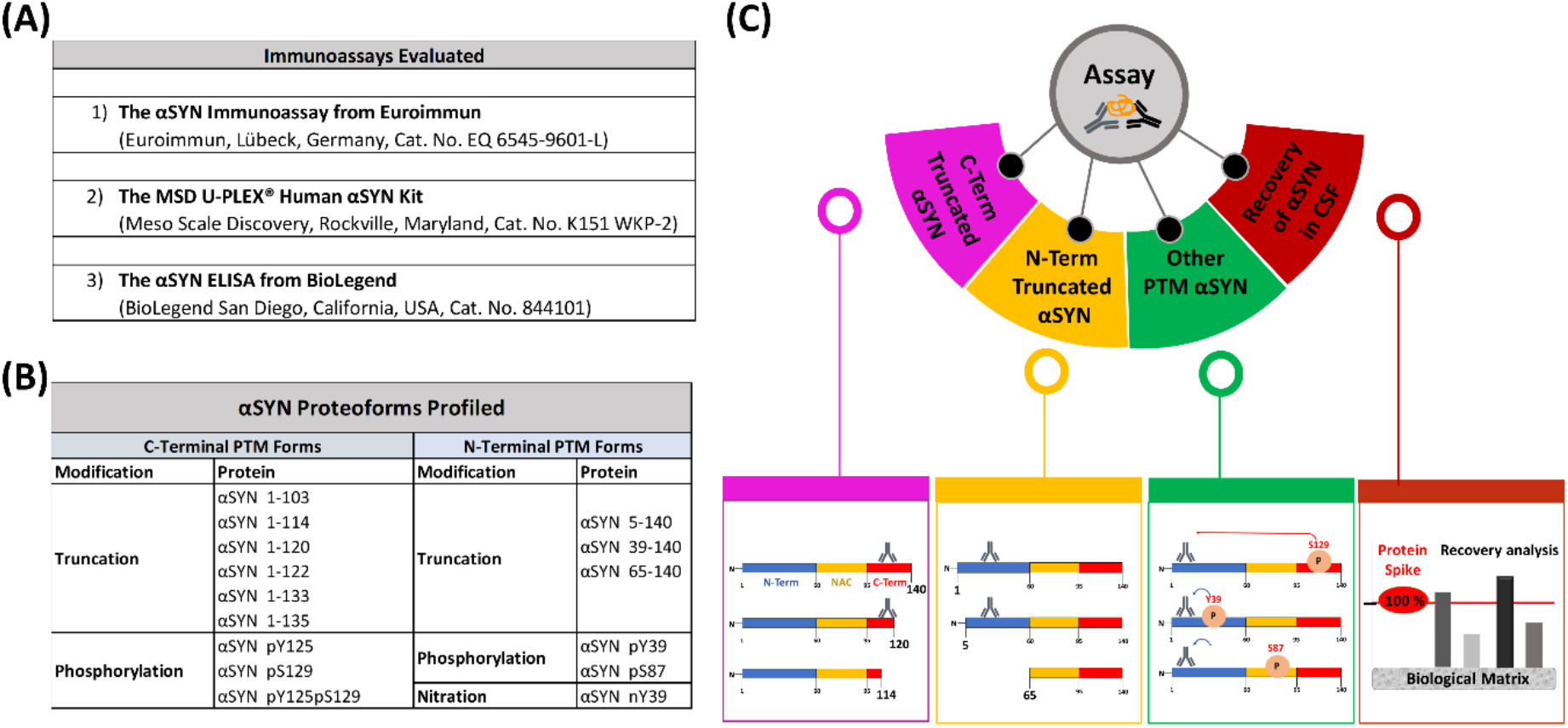
Tables of the evaluated immunoassays (A) and profiled αSYN proteins (B) and (C) schematic depiction of the workflow deployed to evaluate and characterize the immunoassays

### The Euroimmun, Legendmax BioLegend and MSD U-PLEX^®^ immunoassays do not recognize C-terminal truncations ranging from AA 103 to AA 122

A review of the literature on αSYN immunoassays revealed that most antibodies that have been deployed for αSYN detection and measurements target the C-terminus of the protein, spanning residues 103-140 (Figure 1). The situation is similar for the three total αSYN commercial immunoassays used in this study (from Euroimmun, BioLegend, and MSD), with the Euroimmun capture and detection antibodies targeting αSYN 118–125 and αSYN 100–110, respectively, and the Biolegend capture and detection antibodies targeting αSYN 118–122 and αSYN 103–108, respectively. For the MSD immunoassay, only the epitope of the capture antibody (αSYN 110– 125) has been disclosed. The C-terminal region of αSYN is known to harbor several PTMs that occur at multiple residues, including serine (at S129) and tyrosine phosphorylation (Y125, S129, Y133, and Y136), nitration (Y125/Y133/Y136), and truncations (at 103, 114, 119, 120, 122, 133, and 135) [38], [39]. Therefore, we first sought to assess the ability of the three immunoassays to detect C-terminally truncated variants of αSYN cleaved at 103, 114, 120, 122, 133, and 135. We reasoned that this would provide early information on the putative epitopes of the antibodies deployed in these assays and guide the next set of experiments. As such, the specificity of the three assays toward a library of truncated αSYN proteins was analyzed through full dilution curves within the same dynamic range of the kit standard curves obtained by serial dilutions. The results were then fitted to a 4-parameter sigmoid curve and plotted against the kit standard curve. As shown in Figure 3, C-terminally truncated variants of αSYN at amino acid (AA) 103, AA 114, AA 120 and AA 122 were not recognized by any of the three assays (Figure 3 and Supplemental Figures 2-4). In contrast, C-terminally truncated αSYN variants at AA 133 and AA 135 were well detected by all three assays, which is in line with the reported epitopes of some of the antibodies deployed as capturers and detectors. These observations suggest that the presence of truncated αSYN species in biological fluids of interest would not be detected by these antibodies, causing underestimation of the levels of total αSYN by these assays. Furthermore, these findings suggest that the presence of physiological and disease-related C-terminal PTMs that occur in close proximity to the epitopes of the antibodies is likely to interfere with antibody binding and detection of αSYN.

**Figure 3.**
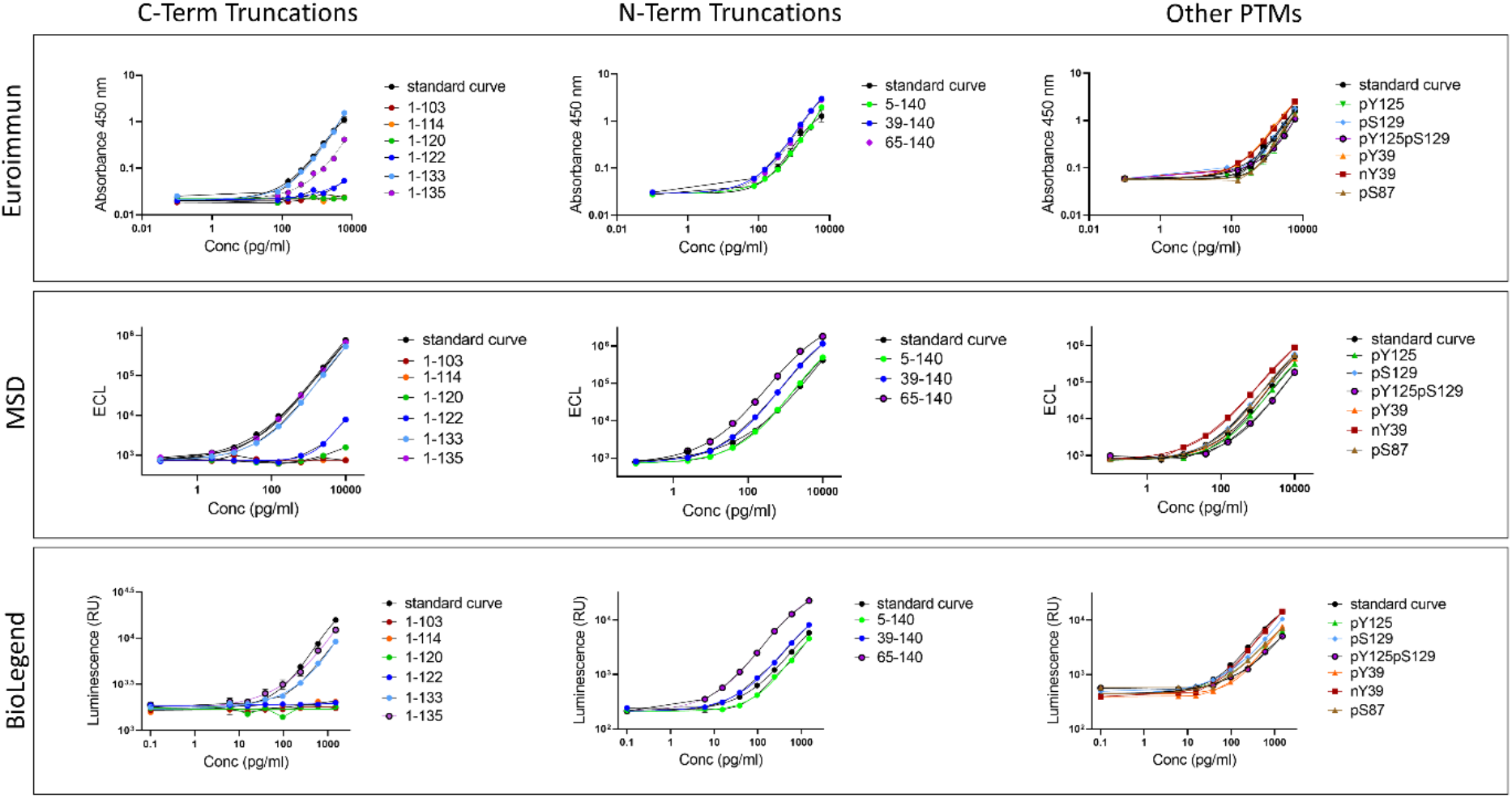
The three assessed immunoassays do not capture the totality of the relevant αSYN species. Full dilution curves of the proteins with the same dynamic range as the standard curve fitted to a 4-parameter sigmoid curve and plotted against the kit standard curve. Data obtained from each assay are detailed separately. C-terminal truncations ranging from 103 to 122 were not recognized by any of the three analyzed assays.

### Several N-terminal truncations and phosphorylation/nitration PTMs differentially influence the detection of αSYN by different immunoassays

Next, we evaluated the ability of the three assays to detect N-terminally truncated forms of αSYN (5-140, 39-140 and 65-140), as well as different forms of nitrated and phosphorylated full length αSYN (nY39, pY39, pS87, pY125, pS129, and pY125/pS129), all of which have been detected in pathological and physiological states in the brain and other tissues [38,39][40] [41]. As expected, all tested N-terminal truncations distant from the disclosed epitopes were well detected by the three kits. Surprisingly, the αSYN protein truncated at position 65 was recognized better than the full-length (FL) standards by all three assays (Figure 3 and Supplemental Figures 2-4). This suggests that certain N-terminal modifications, although distant from the epitopes of the capture and detection antibodies, could induce changes in the conformation of the protein and/or increase epitope exposure/availability compared to the FL standard protein.

Similarly, although phosphorylation and nitration, in general, did not seem to have a major effect on αSYN detection by the different assays, some modifications were recognized slightly better than the FL protein by the Euroimmun and MSD assays (pY39 and nY39 by the Euroimmun assay, and nY39 by the MSD assay), and the Biolegend assay showed less detection of the pY39-, pY125-, and pY125/pS129-modified proteins than the FL αSYN kit standard (Figure 3 and Supplemental Figures 2-4). These observations suggest that the presence of these PTMs in biological tissues or fluids of interest could lead to overestimation of total αSYN levels by the Euroimmun and MSD assays and underestimation of the same total αSYN levels by the Biolegend assay.

### A human CSF matrix effect is observed for most of the analyzed αSYN proteoforms in all three tested immunoassays

To investigate a potential matrix effect of human CSF (hCSF) in the different immunoassays, a selection of 10 PTM αSYN proteins was subjected to spike recovery experiments by spiking assay buffer (AB) and commercial hCSF with high, medium, and low amounts of each protein, according to the dynamic range of the kit’s standard curve. The high, medium and low spikes for the Euroimmun assay were 4000 pg/ml, 1535 pg/ml and 384 pg/ml, 7000 pg/ml, 1000 pg/ml and 100 pg/ml for the MSD assay and 900 pg/ml, 300 pg/ml and 50 pg/ml for the Biolegend assay, respectively. Recovery was assessed as the percentage of the back-calculated values of the spiked samples (with the unspiked sample subtracted) relative to the quantity of nominal spiked protein. High, medium and low recovery percentage were then averaged, and recoveries falling within 80-120% were considered acceptable. The single interpolated values of the three spikes to the standard curve are shown in Supplemental Figures 5-7, and the percentage recovery rates were determined by averaging the rates obtained at high, medium, and low protein spikes across the two matrices (AB and CSF) and are presented in Figure 4A-B. Importantly, this analysis shows that there are indeed differences in how some of the proteins are recovered from the two different matrices. For some of the proteins, the recoveries in AB and hCSF are aligned and mostly reflect the differences previously seen in comparison to the full titration curve of the protein (shown in Figure 3). For example, whereas C-terminally truncated αSYN at 114 (the full titration curve of which was previously not detected) was not recovered in AB or hCSF, N-terminally truncated αSYN at 65 (the full titration curve of which previously showed higher detection than the FL standard) showed high recovery rates in both matrices by all three assays.

**Figure 4.**
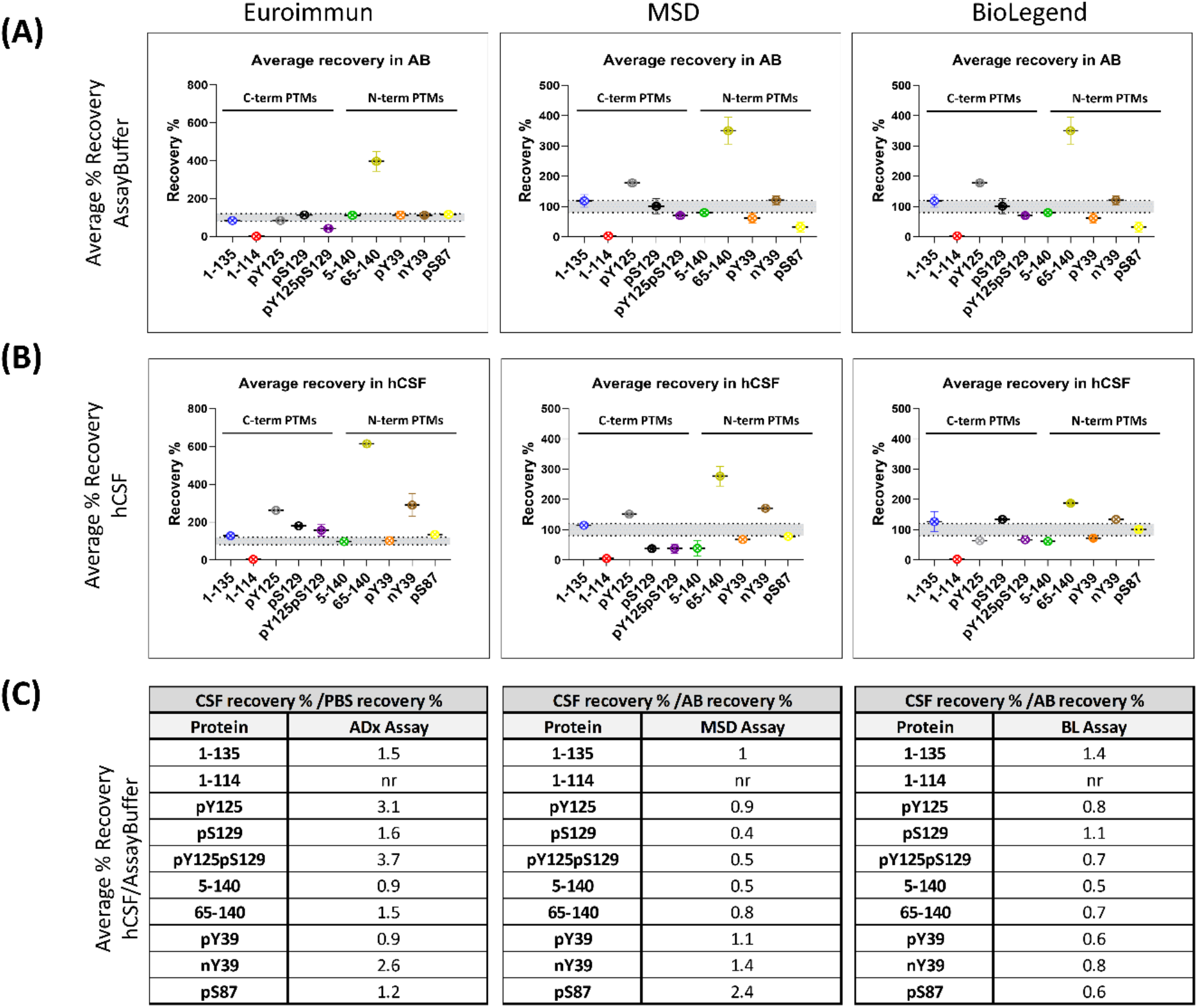
Most PTM αSYN proteins are not fully recovered from human CSF. Recovery is assessed as the percentage of the back-calculated values of the spiked samples subtracted from the unspiked sample with respect to the nominal spiked protein amount. The high, medium and low recovery percentage are averaged. The data obtained from each assay are detailed separately. (A) Average % recovery of spiked PTM αSYN proteins in assay buffer and (B) a commercial pool of human CSF. C) Tables listing the ratios of the average percentage recovery of hCSF to the percentage recovery of AB. Ratio = 1 indicates no difference in recovery from the two matrices. Ratio > 1< indicates a matrix effect.

Notably, other proteins showed different matrix effects in AB and hCSF across the different assays. For instance, phosphorylation at S129 showed high recovery rates with the Euroimmun and Biolegend assays in hCSF but not AB and lower recoveries in hCSF with the MSD assay. Furthermore, phosphorylation at Y39 (pY39), but not nitration on the same residue (nY39), was fully recovered with the Euroimmun assay from AB and hCSF but showed lower recovery in both matrices with the MSD and Biolegend assays. On the other hand, nitration at Y39 (nY39) displays high recovery rates with all the analyzed assays in hCSF but not in AB. Indeed, for all 3 assays, different recovery rates and different matrix effects were observed for pY39 vs. nY39 αSYN. Interestingly, 1-135 αSYN was the only protein that was mostly recovered from both matrices by all three assays, suggesting little matrix effect on this specific proteoform.

To better understand the extent to which different αSYN species are recovered in hCSF compared to AB, we determined the fold-change of the percentage recovery in hCSF over the percentage recovery in AB (Figure 4C). When hCSF recovery rates are divided by the recovery rates in AB, the resulting fold increase/decrease indicates how much better/worse the protein is recovered from the hCSF compared to the AB and, consequently, how strong the matrix effect is for the given protein in the given assay. Ratios of approximately 1 indicate that the recovery in hCSF is approximately the same as that in AB; therefore, there is no matrix effect for the given protein in the given assay. This analysis revealed that most of the proteins show a different extent of fold-change across the two matrices, which is most notable for the pY125, pY125pS129, 65-140, and nY39 proteins; for the nY39, pS87, pS129, pY125pS129, 5-140, and 65-140 proteins; and for the 1-135, pY125pS129, 5-140, 65-140, pY39, and pS87 proteins using the Euroimmun, MSD and Biolegend assays, respectively (Figure 4C). These differences may be caused by the dilution of components in solution that inhibit or enhance the detection of certain proteoforms in the assay method. Lower recoveries in hCSF than in AB for some αSYN proteins could indicate that these species are picked up less due to a matrix effect. It cannot be excluded that endogenous proteases might act on these proteins themselves (as no inhibitors were added to the hCSF) [37,53,54] or that the species might be sequestered in interactions with other components of the matrix. On the other hand, proteins showing higher recoveries in hCSF than in AB either could be endogenously expressed in the matrix and recovered together with the spiked amount or could undergo significant conformational changes that alter their interactions with the antibodies. Altogether, these results demonstrate that none of the three analyzed assays is completely free of hCSF matrix effects on the totality of the analyzed proteins. Furthermore, our findings underscore the importance of validating and/or assessing the suitability of assays, procedures, and antibodies to accurately detect the diversity of αSYN species in different types of samples (e.g., CSF, plasma, saliva, or tissue homogenates).

## Discussion

In this work, we used a library that reconstitutes the diversity of different αSYN proteoforms to determine to what extent three of the most commonly used commercially available total αSYN immunoassays can capture the diversity of αSYN species and provide accurate measurements of total αSYN in biological samples. The library did not cover differences between α, β and γ-synuclein nor αSYN isoforms (140, 126 and 112 kD) but focused on truncated forms and single PTMs of full length αSYN. Our results demonstrate that all three assays failed to recognize C-terminal truncations from AA 103 to AA 122. These findings are in line with the disclosed epitopes of two of the assays. The MSD detection antibody epitope is not disclosed, but our results indicate that it is most likely similar to the those of the other two assays. Moreover, our results demonstrate that truncated αSYN species with cleavages that occur within or near the epitopes of the antibodies used in these assays will not be detected by any of the three assays, leading to underestimation of the total αSYN levels in biological samples.

These findings have significant implications since several C-terminal truncated forms of αSYN (including 1-101, 1-103, 1-115,1-122, 1-124, 1-135 and 1-139) have been found in the brains of PD patients [39,55], and truncations starting at 103 or 119-120 produced by aberrant proteolysis are among the most abundant modified forms of αSYN detected in pathological aggregates [56]. Furthermore, previous immunohistochemical studies and more recent STED microscopy studies demonstrate that C-terminally truncated αSYN accumulates within LBs and is preferentially localized within their cores [57,58]. Increasing evidence also indicates that post-aggregation C-terminal cleavage is an essential step that promotes the efficient packaging of fibrils within LB-like inclusions [59] [48].

Importantly, in addition to their prominent role in pathology initiation and progression, C-terminal truncations have also been proposed to occur under physiological conditions [13,59]. For instance, soluble fractions of aged brains that have not been diagnosed with any type of synucleinopathy show certain truncated αSYN species, particularly 1–119, albeit to a much lesser extent than insoluble fractions from patient brains. It has been postulated that these truncated species are produced by proteases during the physiological clearance processes of endogenous αSYN, including autophagic or lysosomal degradation pathways [59]. Altogether, these findings indicate that truncated αSYN species are prevalent under both physiological and pathogenic conditions and should be considered when developing assays to quantify total αSYN in different types of biological samples. Failure to detect these species will result in significant underestimation of αSYN levels and may contribute to the large variation in the levels of αSYN reported by different groups. It is important to emphasize that the level of C-terminal-truncated αSYN species in biological fluids has not been investigated, and there are no existing assays that allow the specific detection and quantification of these species.

N-terminal truncations at residues 5-140, 39-140 and 65-140 have also been found in the brains of PD patients [39,55]. A recent study of αSYN in the human appendix showed the presence of a mixture of αSYN species that are cleaved at the N- terminus only and αSYN species that are cleaved at both the N- and C-terminal domains of the protein [33]. Herein, we showed that all three assays recognize the three major N-terminally truncated species found in the brain. However, truncation at AA 65 is consistently recognized better than the FL αSYN kit standard by the assays.

Since monomeric αSYN exists as an ensemble of disordered conformations that are stabilized by long-range interactions between the C-terminal domain and the NAC region and N-terminal domain [60], disruption of these interactions can lead to increased exposure of these domains and enhanced antibody binding [61,62]. Indeed, we observed that pY39 and nY39 were recognized better than the standard protein by the Euroimmun assay and nY39 by the MSD assay, while pY39-, pY125-, and pY125/pS129-modified proteins were detected less effectively with the Biolegend assay (Figure 3 and Summarized in Figures 2–4). Therefore, it is plausible to assume that some of the PTMs (such as nitration or phosphorylation at AA 39 or truncation at AA 64) disrupt the long-range interactions between the N- and C-termini of αSYN, leading to better epitope exposure. The possibility that these modifications could also significantly alter the overall conformation of the protein, beyond the modification of N- and C-termini long-range interactions, likewise cannot be excluded.

The relative abundance of some of the αSYN PTMs we studied in human biological fluids has not yet been determined. Nevertheless, one could speculate that the high recovery rates in the hCSF matrix might indicate that some of the modified proteins are endogenously present and recovered along with the spiked protein. This has significant implications for the accuracy of immunoassays that detect certain PTM species better than the FL, as this effect would lead to significant overestimation of the total αSYN levels in biological samples. To the best of our knowledge, N-terminal variants of αSYN have never been used as standards or tools to validate αSYN antibodies and immunoassays.

For the three analyzed assays, we observed that human CSF exerts a matrix effect on several αSYN proteins, as shown by changes in the relative recovery. When the recoveries in hCSF were normalized to the recoveries in AB, the results diverged considerably among the 3 assays. Some proteins were recovered better/worse in hCSF than in AB, and the identity of these proteins was not the same for all 3 assays. Interestingly, phosphorylation at S129 (pS129) was fully recovered in AB by all three analyzed assays but showed high recovery in hCSF with the Euroimmun assay and the Biolegend assays and low recovery in hCSF with the MDS assay. For the Euroimmun and Biolegend assays, only 4 out of the 10 proteins analyzed were not influenced by the hCSF matrix, while for the MSD assay, 5 out of 10 did not show any matrix effect. As the identity of these matrix-insensitive proteins is different across the three assays, measurements of “total αSYN levels” of the same hCSF sample containing PTMs using these three assays would differ considerably and would provide a solid explanation for the previously observed large variation in the absolute concentrations of total αSYN quantified by these three assays using the same CSF samples [37]. Notably, a similar impact of the CSF matrix on the detection and quantitation of analytes has been previously reported for amyloid-beta [63], [64], where the matrix affecting the measurements is discussed as a possible contributor to the low diagnostic power of these assays.

Although the capture antibodies used in the three immunoassays target similar epitopes in the C-terminal domain of αSYN, the assays exhibit significantly different sensitivities to different αSYN species and show different matrix effects, all of which impact homogenous protein recovery in hCSF. Other factors contributing to these differences include whether and how the antibody is immobilized on the surface or if the analyte recognition is carried out in solution. The sensitivity of the platform itself could also account for differences, as low-abundance species might contribute to the total signal in a different way when an ultrasensitive platform is used. Less sensitive platforms might not be able to measure low-abundance species, therefore overlooking and *de facto* flattening the contribution of these proteins to total αSYN levels, compared to that obtained by ultrasensitive platforms with a much lower limit of detection. For instance, the increased αSYN levels in plasma from PD patients observed with the ultrasensitive Quanterix assay compared to other less sensitive methods could be related to the increased sensitivity of this platform, which may facilitate the detection and quantitation of low-abundance forms of αSYN, as discussed recently in [21]. On the other hand, assays with low dynamic ranges of their standard curves and low upper limits of quantification (ULOQ) will have decreased sensitivity and might account for saturation of the signal when certain species are highly abundant in the analyzed matrix. Accordingly, in both cases, these assays would not provide a reliable measurement of total αSYN levels.

Altogether, our results show that the three immunoassays (Euroimmun, MSD, and Biolegend) do not capture the totality of the relevant αSYN species and therefore may not be appropriate tools to provide an accurate estimate of total αSYN levels, at least until the distribution and levels of different αSYN-modified proteins in different biological samples are accurately determined. Moreover, we have identified a matrix effect of hCSF that differs across different proteins and immunoassays. It cannot be ascribed only to the different antibody epitopes, as they are shared among the assays. Our work highlights the critical importance of pre-assay development including the thorough validation of deployed antibodies and assays using well-characterized αSYN proteins, as this will dictate the downstream specificity of the immunoassay and ensure that the antibodies deployed accurately capture the diversity and complexity of αSYN proteoforms in biological fluids. Previous studies attributed discrepancies in the absolute values of total αSYN measured by different research teams to different factors, including heterogeneity of patient populations, differences in sample preparation and handling procedures, matrix effects, and the use of poorly characterized protein standards. Our work here supports some of these findings and highlights that insufficient pre-assay validation of antibodies and lack of validation of the assays using standards that reproduce the complexity of the αSYN proteome are two other major contributing factors.

The development of a robust and quantitative total αSYN assay is urgently needed. It would enable the discovery of novel biomarkers based on measuring the ratios of specific forms of αSYN to total αSYN. As most PTMs represent a small percentage of total αSYN, depending on the sensitivity of the detection method used, the ratio of one PTM to the total αSYN levels could be more informative and might better correlate with the disease than the levels of total αSYN or individual PTMs alone. Indeed, all existing commercially available assays were developed to target predominantly monomeric and/or unmodified aggregated forms of αSYN [65]. Given the tight association between αSYN PTMs and αSYN aggregation and pathology initiation, the development and validation of future assays must incorporate the need to capture and detect modified aggregated forms of the protein. The development of immunoassays capable of capturing and measuring the complexity of αSYN in human biological fluids would enable more systematic studies to determine if levels of total αSYN or certain αSYN proteoforms could serve as useful diagnostic biomarkers for early PD diagnosis, monitoring of disease progression and assessment of target engagement for novel αSYN-targeting therapies. The identification of capture and detection antibodies that detect all of the relevant αSYN species, including pathogenic conformations and PTM forms of the protein, would represent an important step toward achieving this goal.

## Acknowledgments

This work was supported by the Michael J. Fox Foundation under Research Grants MJFF-15995 and MJFF-007894 and the EPFL. We are grateful to Peggy Taylor, Jesse Ni and the Neuroscience team at Biolegend and to Erik Stoops and his team at ADx NeuroSciences NV for the kit supply, for independently validating some of our results, and for their support and assistance throughout the project.

## Author’s contribution

HAL conceived and conceptualized the study. HAL, MBF, and LP designed the experiments. LHE and LA performed preliminary pilot experiments. LP coordinated, supervised, and led the implementation of all the experiments in the manuscript and analyzed the data with the help of NC. RB and ES contributed valuable reagents for the project. HAL, LP, and MBF wrote the manuscript. All the authors reviewed the final version and approved its submission.

## Conflict of interest

Hilal Lashuel (HAL) has received funding from industry to support research on neurodegenerative diseases, including from Merck Serono, UCB, Idorsia and Abbvie. These companies had no specific role in the in the conceptualization and preparation of and decision to publish this work. HAL is also the Co-founder and Chief Scientific Officer of ND BioSciences SA, a company that develops diagnostics and treatments for neurodegenerative diseases based on platforms that reproduce the complexity and diversity of proteins implicated in neurodegenerative diseases and their pathologies.

Mohamed Bilal Fares (MBF) is the Co-founder and Director of R&D of ND BioSciences SA.

Lara Petricca (LP) is the Head of Biomarkers and Diagnostic Development of ND BioSciences SA.

Erik Stoops is an employee and shareholder of ADx NeuroSciences.

