## Supplementary data for "Comparative analysis of total alpha-synuclein (αSYN) immunoassays reveals that they do not capture the diversity of modified αSYN proteoforms"

1 ND Biosciences SA, Epalinges, Switzerland.

2 Laboratory of Molecular and Chemical Biology of Neurodegeneration, Brain Mind Institute, Ecole Polytechnique Fédérale de Lausanne (EPFL), BMI SV LMNN Station 19, 1015 CH Lausanne, Switzerland.

3 ADx NeuroSciences NV, Technologiepark 94 – Bio Incubator, 9052 Gent, Belgium

Laboratory of Molecular and Chemical Biology of Neurodegeneration, EPFL SV BMI LMNN, AI 2151 (Bâtiment AI), Station 19, CH-1015 Lausanne, +41216939691

Running title: Comparison of alpha-synuclein immunoassays

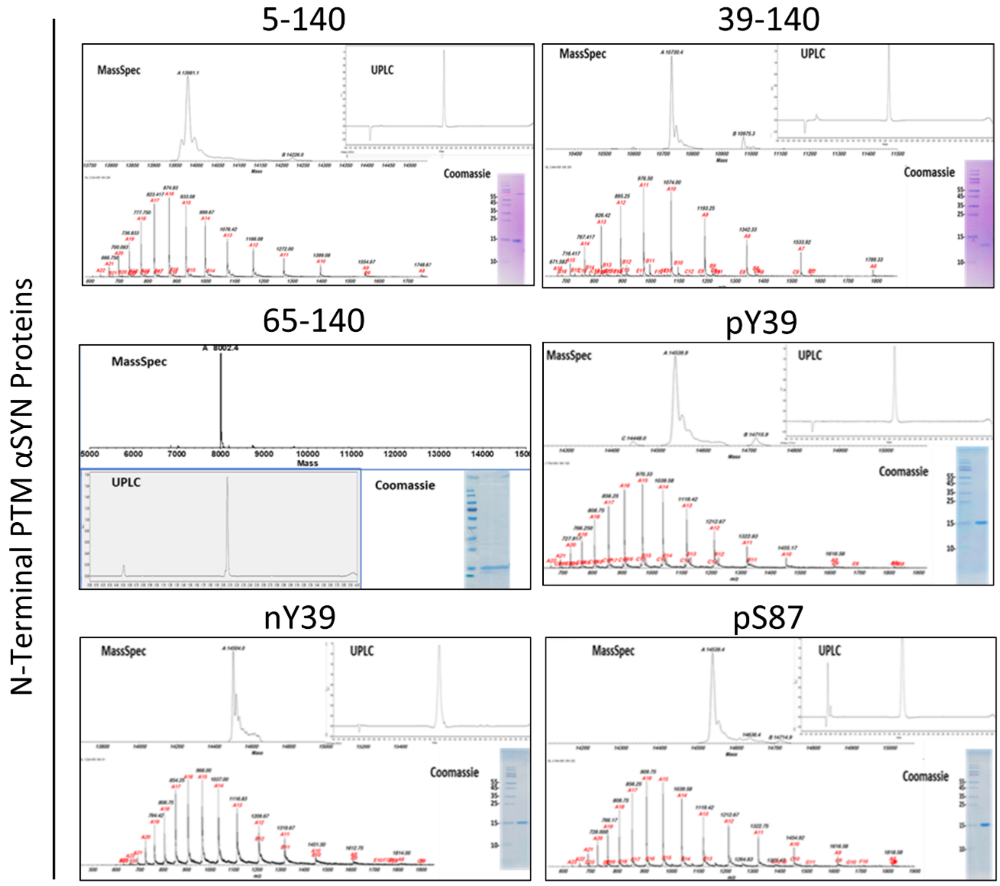


| N-Terminal PTM Human αSYN Proteins | |
|---|---|
| **Protein** | **Molecular Weight (Da)** |
| αSYN 5-140 | 13967.524 |
| αSYN 39-140 | 10599.583 |
| αSYN 64-140 | 8103.7896 |
| αSYN pY39 | 14540.094 |
| αSYN nY39 | 14505.000 |
| αSYN pS87 | 14540.094 |

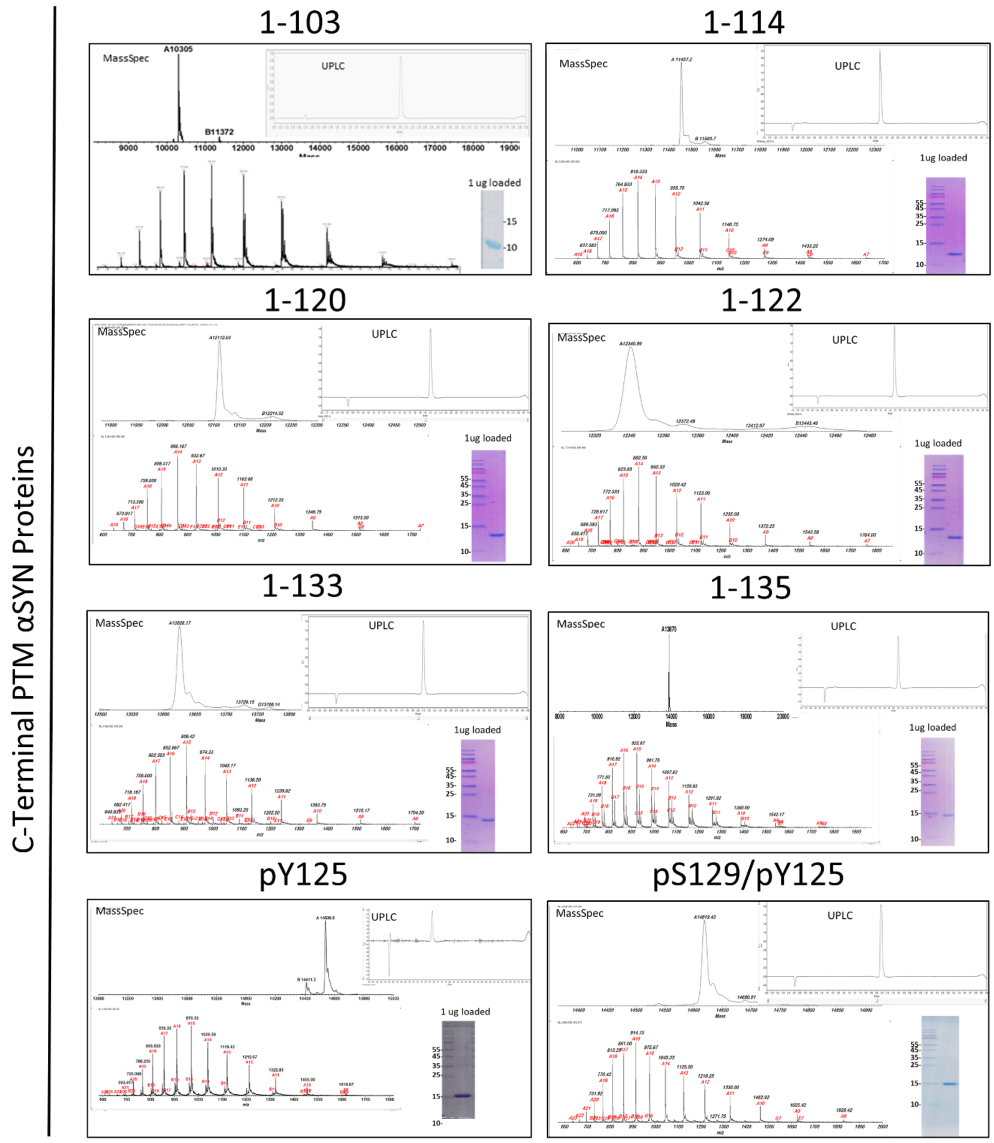


| C-Terminal PTM Human αSYN Proteins | |
|---|---|
| **Protein** | **Molecular Weight (Da)** |
| αSYN 1-103 | 10303.833 |
| αSYN 1-114 | 11457.040 |
| αSYN 1-120 | 12111.774 |
| αSYN 1-122 | 12340.966 |
| αSYN 1-133 | 13627.294 |
| αSYN 1-135 | 13870.513 |
| αSYN pY125 | 14540.000 |
| αSYN pY125 pS129 | 14620.074 |

**Figure S1. Characterization of the generated library of αSYN proteins bearing the most commonly occurring N-terminal and C-terminal PTMs.**

Mass spectrometry, ultra-performance liquid chromatography (UPLC) and Coomassie staining were performed to establish the purity and integrity of all generated proteins.

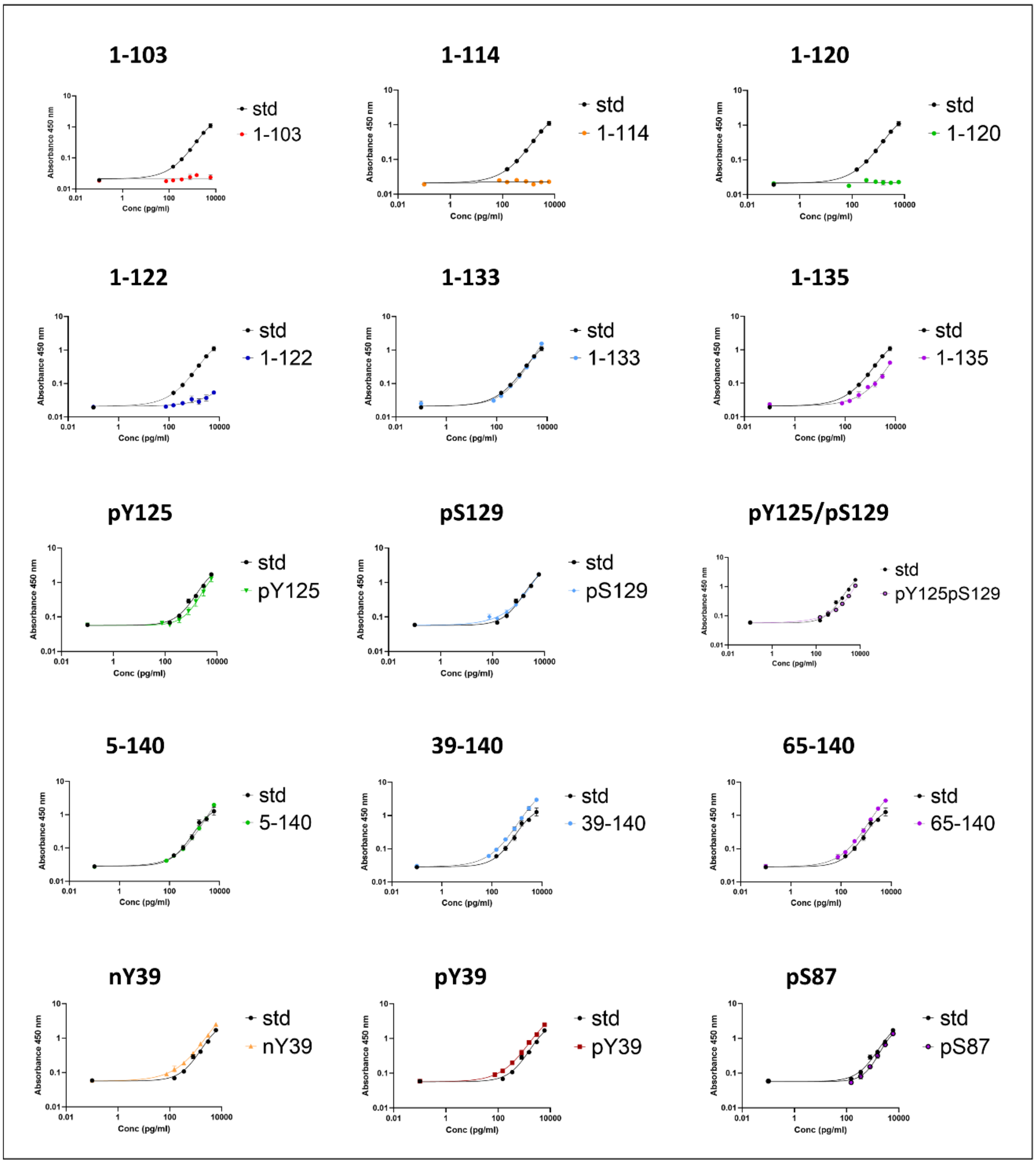


**Figure S2. Single graphs separately showing the detection of PTM αSYN proteins using the Euroimmun immunoassay.**

Full dilution curves of the proteins with the same dynamic range as the standard curve fitted to a 4-parameter sigmoid curve and plotted against the kit standard curve. C-terminal truncations ranging from 103 to 122 were not recognized by any of the analyzed assays.

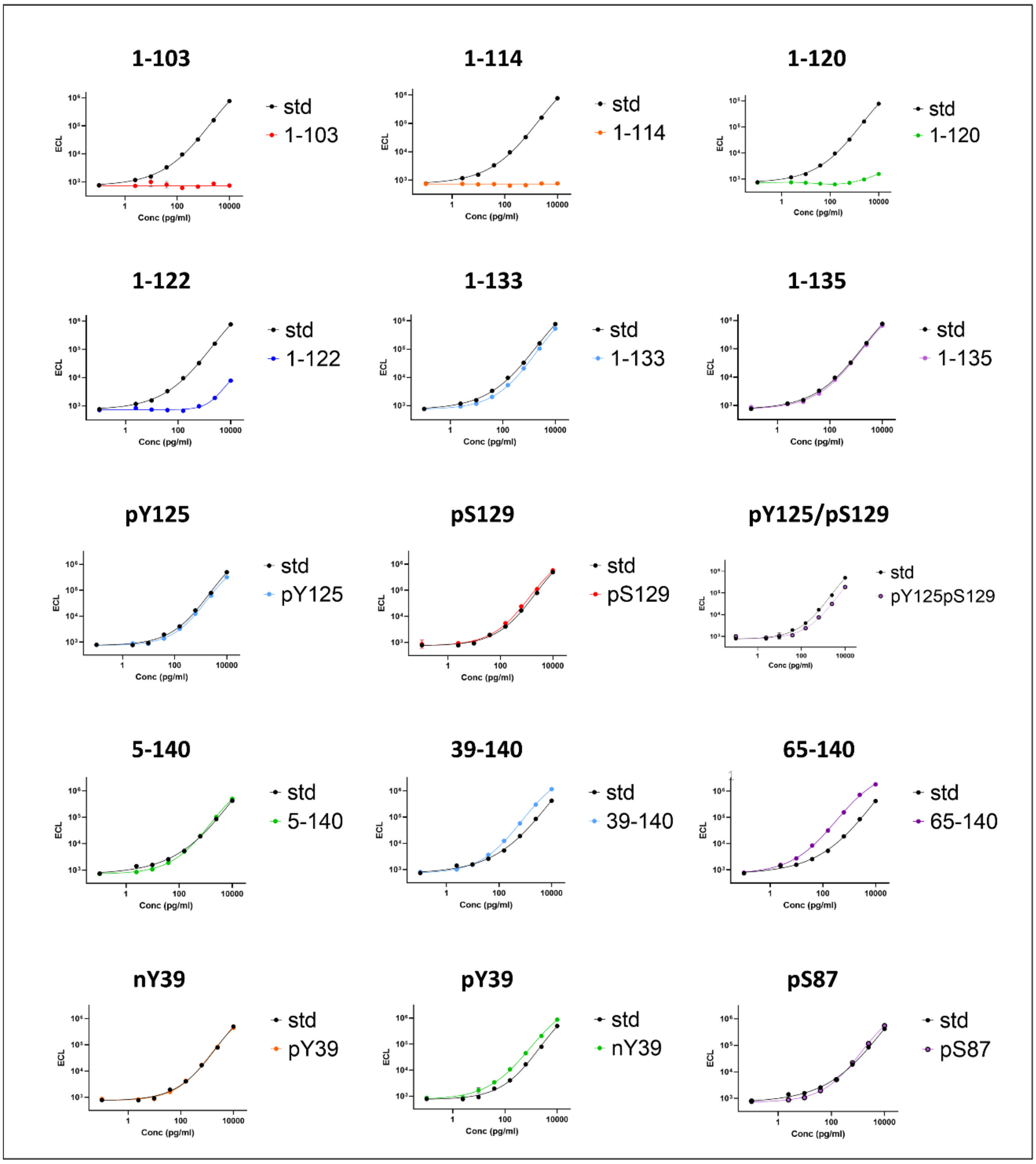


**Figure S3. Single graphs separately showing the detection of PTM αSYN proteins using the MSD immunoassay.**

Full dilution curves of the proteins with the same dynamic range as the standard curve fitted to a 4-parameter sigmoid curve and plotted against the kit standard curve. C-terminal truncations ranging from 103 to 122 were not recognized by any of the analyzed assays.

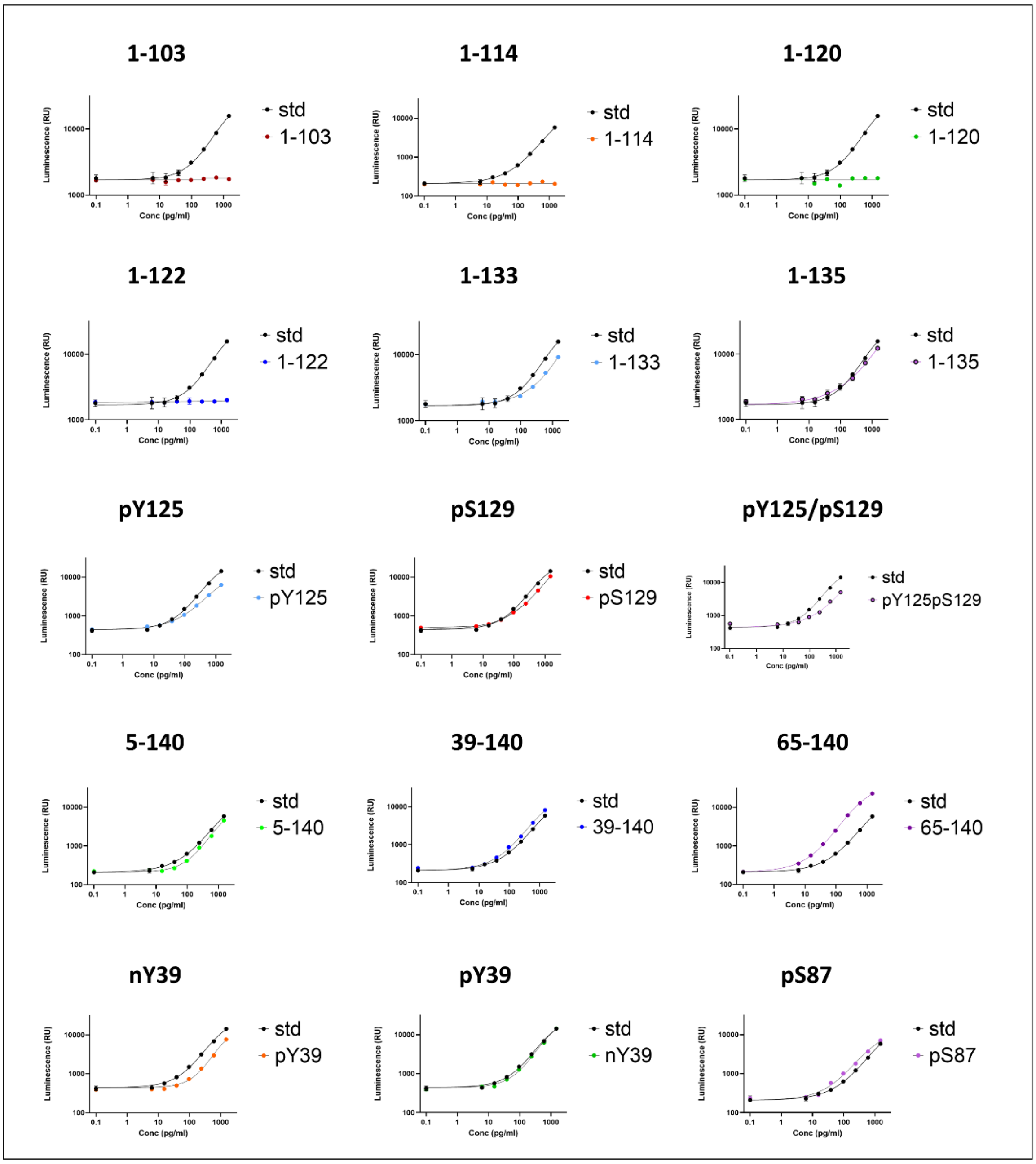


**Figure S4. Single graphs separately showing the detection of PTM αSYN proteins using the Biolegend immunoassay.**

Full dilution curves of the proteins with the same dynamic range as the standard curve fitted to a 4-parameter sigmoid curve and plotted against the kit standard curve. C-terminal truncations ranging from 103 to 122 were not recognized by any of the analyzed assays.

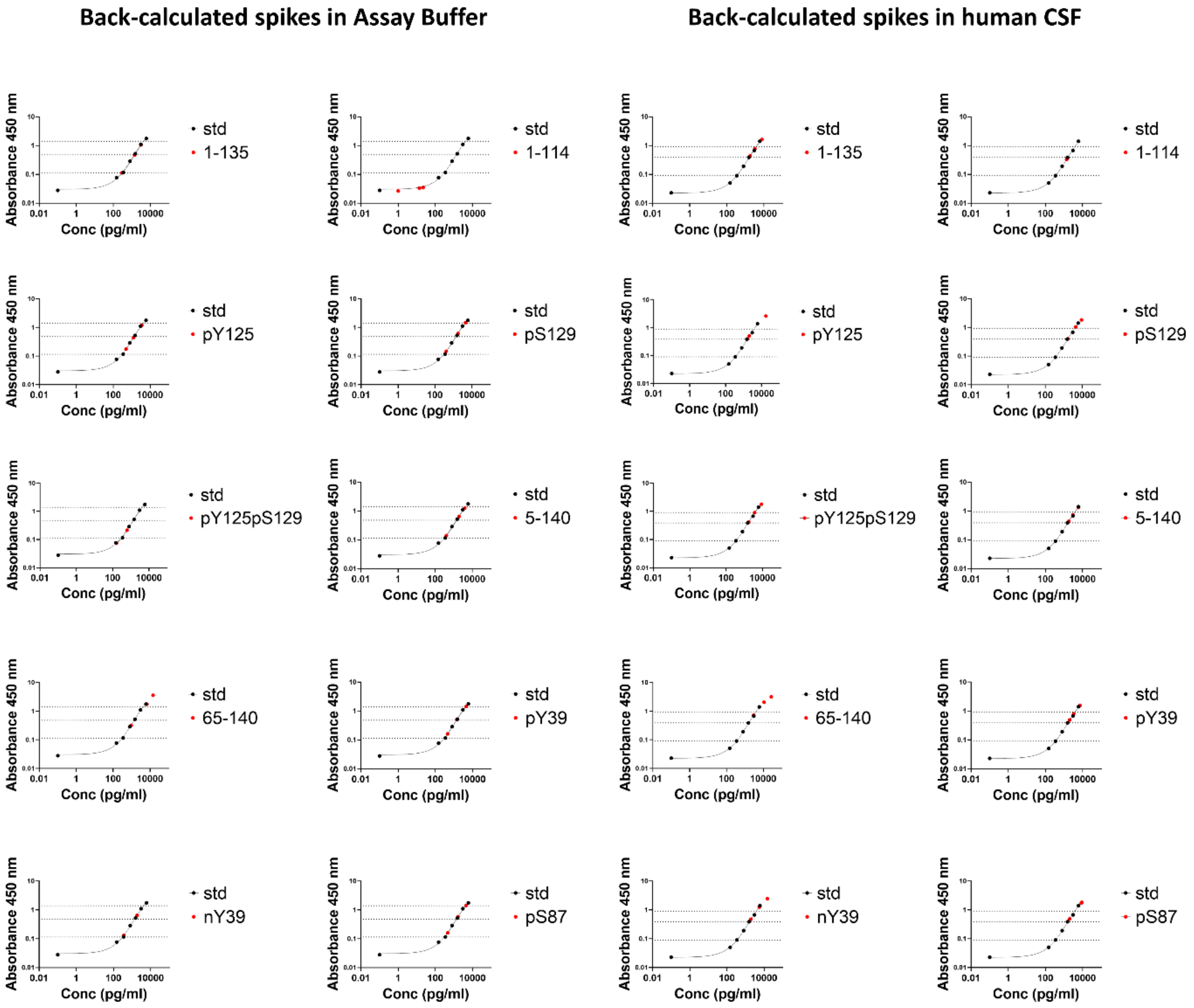


**Figure S5. Euroimmun immunoassay.**

Single graphs separately showing the back-calculated values of the high-, medium- and low-PTM αSYN spikes in AB interpolated to the assay standard curve (red dots). Nominal expected high, medium and low spikes interpolated to the standard curve are shown by the dotted lines.

**Back-calculated spikes in Assay Buffer** **Back-calculated spikes in human CSF**

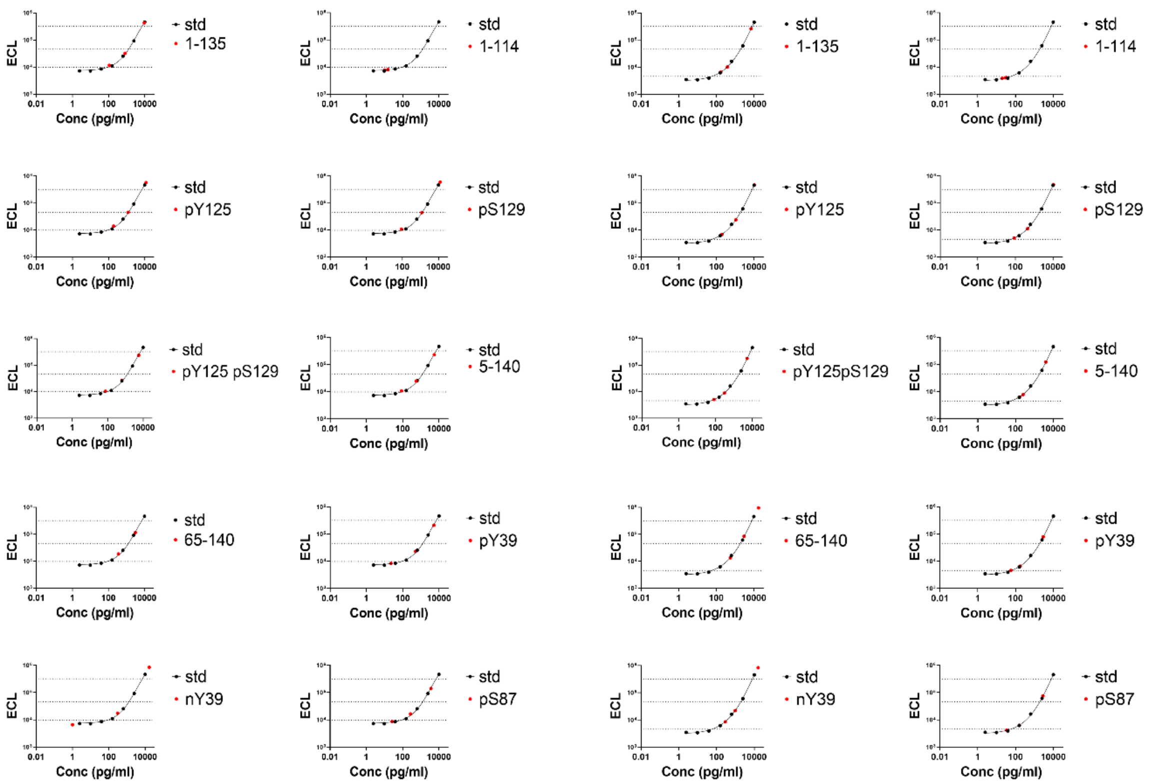


**Figure S6. MSD immunoassay.**

Single graphs separately showing the back-calculated values of the high-, medium- and low-PTM αSYN spikes in AB interpolated to the assay standard curve (red dots). Nominal expected high, medium and low spikes interpolated to the standard curve are shown by the dotted lines.

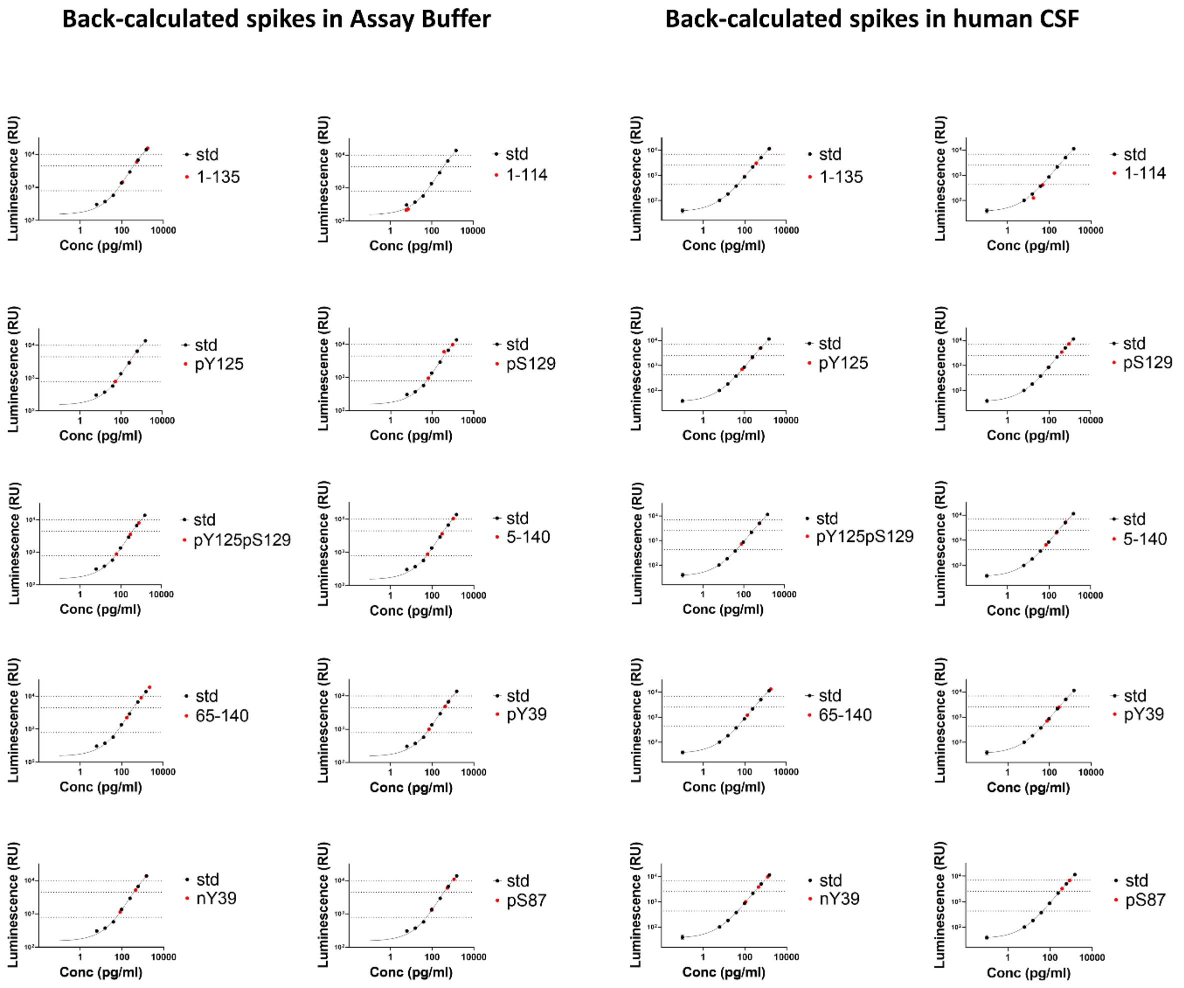


**Figure S7. Biolegend immunoassay.**

Single graphs separately showing the back-calculated values of the high-, medium- and low-PTM αSYN spikes in AB interpolated to the assay standard curve (red dots). Nominal expected high, medium and low spikes interpolated to the standard curve are shown by the dotted lines.

| Evaluated parameter | Euroimmun | MSD | Biolegend |
|---|---|---|---|
| Well Format | 96 | 96 | 96 |
| Readout | Absorbance 450 nm | ECL | Luminescence |
| Dynamic Range pg/ml | 5988-150 | 10'000-2,44 | 1500-6,1 |
| Internal Calibrator controls | ✓ | can be purchased | ✗ |
| Easy to handle | ✓ | ✓ | ✗ |
| Assay running time | 5 hrs over 1 day | 4 hrs over 1 day | 2 days |
| Final reaction stability | for 1 hr | ✗ | ✗ |
| Reading of partial plate | ✓ | ✓ | ✗ |

**Figure S8. Summary table of general technical and handling considerations of the analyzed total αSYN immunoassay kits.**
